# Emergence and Genetic Analysis of Avian Gyrovirus 2 Variants-Related *Gyrovirus* in Farmed King-ratsnakes (*Elaphe carinata*): the First Report in China

**DOI:** 10.1101/629980

**Authors:** Qianqian Wu, Qinxi Chen, Wen Hu, Xueyu Wang, Jun Ji

**Author notes:** **Corresponding authors**: Jun Ji, Henan Provincial Engineering Laboratory of Insects Bio-reactor, Wolong Road 1638, Nanyang Normal University, Nanyang, PR China, E-mail addresses.

## Abstract

Avian gyrovirus 2 (AGV2), which is similar to chicken infectious anemia virus, is a new member of the *Circovirus* genus. AGV2 has been detected not only in chicken but also in human tissues and feces. In this study, a total of 91 samples (8 liver tissues and 83 faecal samples) collected from king-ratsnakes (*Elaphe carinata*) at 7 separate farms in Hubei and Henan, China, were analyzed to detect AGV2 DNA via specific PCR. The results indicated a low positive rate of AGV2 (6.59%, 6/91) in the studied animals, and all of the positive samples were from the same farm. The AGV2 strain HB2018S1 was sequenced, and the genome with a total length of 2376 nt contained three partially overlapping open reading frames: VP1, VP2, and VP3. Phylogenetic tree analysis revealed that the HB2018S1, NX1506-1 strain from chickens in China belong to the same clade, with nucleotide homology as high as 99.5%. In total, 10 amino acid mutation sites, including 44R/K, 74T/A, 256 C/R, 279L/Q, and 373V/A in AGV2 VP1; 60I/T, 125T/I, 213D/N, and 215L/S in AGV2 VP2; and 83H/Y in AGV2 VP3, were found in the genome of HB2018S1 that were different from those observed in most reference strains, suggesting that the differences are related to an transboundary movement among hosts which needs to be further elucidated.

**IMPORTANCE:** Recently, AGV2 has been detected in live poultry markets and human blood in mainland China. Previous findings indicated future studies should investigate the large geographic distribution of AGV2 and monitor the variants, the host range, and the associated diseases. To the best of our knowledge, this study is the first report on AGV2 infected poikilotherm, suggested that cross-host transmission of viruses with circular single-stranded DNA genomes would be a public health concern.

## INTRODUCTION

Avian gyrovirus 2 (AGV2) belongs to the viral genus *Cyclovirus* (family: *Circoviridae*), and it was first detected in diseased chicken in Brazil in 2011[1]. AGV2 infections in chicken can result in brain damage, mental retardation, and weight loss[2]. Although other specific symptoms of AGV2 infections have not been confirmed, autopsy-based studies have reported clinical manifestations such as hemorrhage, edema, glandular gastric erosion, and facial and head swelling in infected chickens[3]. Varela et al. (2014) used a duplex quantitative real-time PCR assay to detect commercially available poultry vaccines and suggested that the widespread existence of AGV2 is associated with vaccine contamination [4]. AGV2 has also been reported in different regions of Europe, Latin America, Africa, South America, and Asia, which indicates its global distribution [5-8]. Further, AGV2 has been detected not only in poultry but also in human blood samples [9]. In 2011, Sauvage et al. (2011) found human gyrovirus, which is similar to AGV2, in a skin swab of a healthy person, suggesting that AGV2 has public health significance [10]. In the present study, we detected AGV2 in farmed snakes that was highly homologous to AGV2 and amplified its whole-genome sequence. We subsequently performed an in-depth sequence analysis based on genetic evolution and amino acid mutations between the strain and reference strains.

## RESULTS

### Positive rates of AGV2 in snakes in China

To increase our understanding of the infection of AGV2 in farmed snakes in China, a total of 91 king-ratsnakes (*Elaphe carinata*) samples were collected and tested. The results indicated that 5 faecal samples (6.02%, 5/83) and only one liver tissues (12.5%, 1/8) were positive for AGV2. All the 6 positive samples were collected from the same snake farm in Hubei, China. Detailed information on the snake samples testing positive for AGV2 DNA is listed in Table 1.

**Table 1.** Information regarding the samples of snake in our study

| Farm | Province | Number of samples | Positive rates |  |
| --- | --- | --- | --- | --- |
|  |  |  | Faecal samples | Liver samples |
| A | Henan | 10 | 0/9 | 0/1 |
| B | Hubei | 13 | 5/11 | 1/2 |
| C | Hubei | 5 | 0/5 | 0/0 |
| D | Hubei | 21 | 0/20 | 0/1 |
| E | Henan | 19 | 0/17 | 0/2 |
| F | Hubei | 11 | 0/10 | 1/1 |
| G | Hubei | 12 | 0/11 | 1/1 |
| Total |  | 91 | 5/83 | 1/8 |

### Sequence analysis of whole-genome

Based on the results of sequencing and contig-assembly, the AGV2 strain was named HB2018S1 and was found to contain a whole genome with a full length of 2376 nt. The nucleotide homology of HB2018S1 with the same cluster of the NX1506-1, NX1506-2, NX1510, and JL1508 reference strains was up to 99.5%. The lowest nucleotide homology of 95.3% was detected between HB2018S1 and S53/It from chickens. In addition, the nucleotide homologies between HB2018S1 and each of the three human strains 915F06007FD, CL33, and JQ69076 3 were 95.6%, 97.1%, and 96.8%, respectively.

### Phylogenetic analysis

Phylogenetic tree analysis revealed that AGV2 sequences of the present study and the 25 full-length genome sequences downloaded from GenBank could have two branches (Fig. 1). HB2018S1 and most of the downloaded sequences belonged to branch I, which was subsequently divided into four clusters. Excluding one cluster containing two strains of the human virus 915F06007FD and CL33, and G13 from ferrets, the remaining three clusters were found to represent AGV2 mainly from chickens.

**Figure 1.**
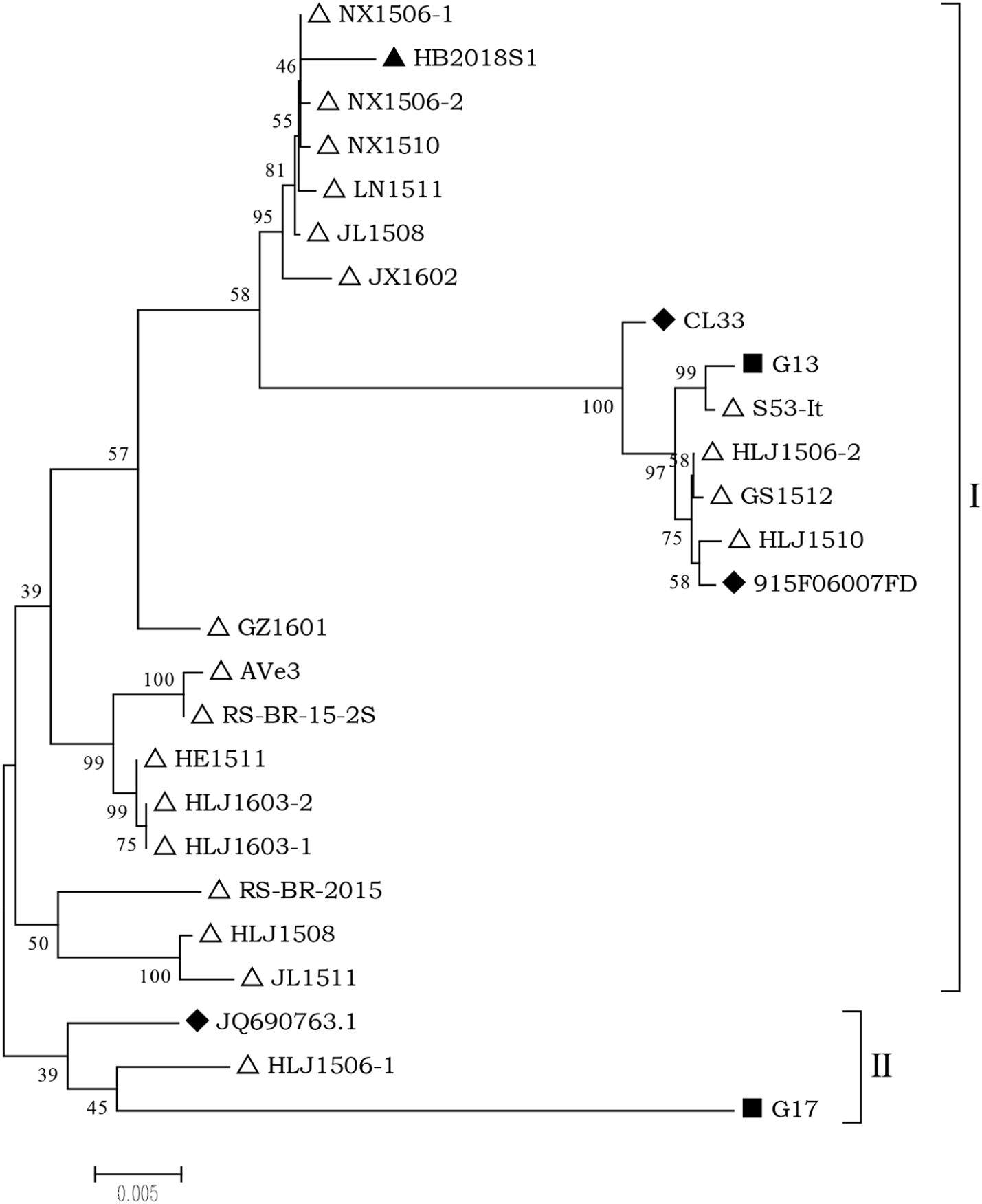
Phylogenetic analysis of the nucleotide sequences of HB2018S1 and 25 reference genomes available at GenBank. “△” indicates that the host of the strain is chicken. “◆” indicates that the host of the strain is human. “■” indicates that the host of the strain is ferret. “▲” indicates the strain from snake in the present study.

### Major mutation analysis of mutated amino acids

There are three overlapping ORFs in the AGV2 genome, which encode VP1, VP2, and VP3 proteins. ORF3, which is located at sites 953–2335 and encodes VP1, comprises 460 amino acid residues. VP1 of HB2018S1 shared 97.4%–98.9% homology of deduced amino acid with the reference strains. VP1 of all strains contained 20 substituted amino acids (displayed in gray in Table 2), including at sites 36, 44, 74, 95, 154, 212, 242, 256, 270, 279, 288, 293, 310, 311, 314, 373, 383, 401, 416, and 459, and VP1 of HB2018S1 had five unique amino acid mutations at sites 44R/K, 74T/A, 256C/R, 279L/Q, and 373V/A.

**Table 2.** Mutations at different amino acid sites in AGV2 VP1

| Strains | 36<br>G/S | 44<br>R/K | 74<br>T/A | 95<br>K/T | 154<br>A/S | 212<br>K/R | 242<br>R/G | 256<br>C/R | 270<br>S/A | 279<br>L/Q | 288<br>V/Q | 293<br>G/Q | 310<br>E/Q | 311<br>D/E | 314<br>R/K | 373<br>V/A | 383<br>P/Q | 401<br>V/M | 416<br>L/I | 459<br>N/T |
| --- | --- | --- | --- | --- | --- | --- | --- | --- | --- | --- | --- | --- | --- | --- | --- | --- | --- | --- | --- | --- |
| HB2018S1 | G | K | A | K | A | K | R | R | S | Q | V | G | E | D | R | A | P | V | L | N |
| NX1510 | G | R | T | K | A | K | R | C | S | L | V | G | E | D | R | V | P | V | L | N |
| NX1506-2 | G | R | T | K | A | K | R | C | S | L | V | G | E | D | R | V | P | V | L | N |
| NX1506-1 | G | R | T | K | A | K | R | C | S | L | V | G | E | D | R | V | P | V | L | T |
| JL1508 | G | R | T | K | A | K | R | C | S | L | V | G | E | D | R | V | P | V | L | N |
| LN1511 | G | R | T | K | A | K | R | C | S | L | V | G | E | D | R | V | P | V | L | T |
| JX1602 | G | R | T | K | A | K | R | C | S | L | V | G | E | D | R | V | P | V | L | N |
| GZ1601 | S | R | T | K | A | K | R | C | S | L | V | G | E | D | R | V | P | V | L | N |
| HLJ1603-2 | G | R | T | K | A | K | R | C | S | L | V | G | E | D | R | V | P | V | L | N |
| HLJ1603-1 | G | R | T | K | A | K | R | C | S | L | V | G | E | D | R | V | P | V | L | T |
| HE1511 | G | R | T | K | A | K | R | C | S | L | V | G | E | D | R | V | P | V | L | T |
| HLJ1506-2 | G | R | T | K | A | K | R | C | S | L | V | G | E | D | R | V | P | V | L | N |
| GS1512 | G | R | T | K | A | K | R | C | S | L | V | G | E | D | K | V | P | V | L | N |
| HLJ1508 | G | R | T | T | S | K | R | C | S | L | Q | Q | E | D | R | V | P | V | L | N |
| G13 | G | R | T | K | A | K | R | C | S | L | V | G | E | E | R | V | P | V | L | N |
| JL1511 | G | R | T | T | S | K | R | C | S | L | Q | Q | E | D | R | V | P | V | L | N |
| HM590588 | G | R | T | T | S | R | G | C | A | L | V | G | Q | D | R | V | Q | M | I | N |
| JQ690763 | G | R | T | K | A | K | R | C | S | L | Q | Q | E | D | R | V | P | V | L | N |
| RS/BR/15/2S | G | R | T | K | S | K | R | C | A | L | V | G | Q | D | R | V | Q | M | I | N |
| HLJ1506-1 | S | R | T | K | A | K | R | C | S | L | V | G | E | D | R | V | P | V | L | N |
| 915F06007FD | G | R | T | K | A | K | R | C | S | L | V | G | E | D | R | V | P | V | L | N |
| G17 | S | R | T | K | S | K | R | C | S | L | Q | Q | E | D | K | V | P | V | L | N |
| CL33 | G | R | T | K | A | K | R | C | S | L | V | G | E | D | R | V | P | V | L | N |
| S53/It | G | R | T | K | A | K | R | C | S | L | V | G | E | D | R | V | P | V | L | N |
| HLJ1510 | G | R | T | K | A | K | R | C | S | L | V | G | E | D | R | V | P | V | L | N |

The ORF1 encoding AGV2 VP2 was located at sites 450–1145 and encoded 231 amino acid sequences, and VP2 sequences of HB2018S1 shared 94.0%–98.3% amino acid homology with those of the reference strains. There were 15 hyper-mutated amino acid sites in all sequences (14N/T, 60I/T, 125T/I, 141R/Q, 156–158GKR/RRG, 161Y/H, 165A/T, 167T/S, 174–175EE/DD, 179A/V, 213D/N, and 215L/S), whereas G17 from ferrets had an S insertion at site 162. In addition, the substitutions 60I/T, 125T/I, 213D/N, and 215L/S only occurred in HB2018S1 (displayed in gray in Table 3).

**Table 3.** Mutations at different amino acid sites in AGV2 VP2

| Strains | 14<br>N/T | 60<br>I/T | 125<br>T/I | 141<br>R/Q | 156-158<br>GKR/RRG | 161<br>Y/H | 162<br>- | 165<br>A/T | 167<br>T/S | 174-175<br>EE/DD | 179<br>A/V | 213<br>D/N | 215<br>L/S |
| --- | --- | --- | --- | --- | --- | --- | --- | --- | --- | --- | --- | --- | --- |
| HB2018S1 | N | T | I | R | GKR | Y | - | A | T | EE | A | N | S |
| NX1510 | N | I | T | R | GKR | Y | - | A | T | EE | A | D | L |
| NX1506-2 | N | I | T | R | GKR | Y | - | A | T | EE | A | D | L |
| NX1506-1 | N | I | T | R | GKR | Y | - | A | T | EE | A | D | L |
| JL1508 | N | I | T | R | GKR | Y | - | A | T | EE | A | D | L |
| LN1511 | N | I | T | R | GKR | Y | - | A | T | EE | A | D | L |
| JX1602 | N | I | T | R | GKR | Y | - | A | T | EE | A | D | L |
| GZ1601 | T | I | T | Q | RRG | H | - | T | T | DD | V | D | L |
| HLJ1603-2 | T | I | T | Q | RRG | H | - | T | T | DD | V | D | L |
| HLJ1603-1 | T | I | T | Q | RRG | H | - | T | T | DD | V | D | L |
| HE1511 | T | I | T | Q | RRG | H | - | T | T | DD | V | D | L |
| HLJ1506-2 | N | I | T | R | GKR | Y | - | A | T | EE | A | D | L |
| GS1512 | N | I | T | R | GKR | Y | - | A | T | EE | A | D | L |
| HLJ1508 | N | I | T | Q | RRG | H | - | A | T | DD | V | D | L |
| G13 | N | I | T | R | GKR | Y | - | A | T | EE | A | D | L |
| JL1511 | N | I | T | Q | RRG | H | - | A | T | DD | V | D | L |
| HM590588 | T | I | T | Q | RRG | H | - | T | T | DD | V | D | L |
| JQ690763 | N | I | T | Q | RRG | H | - | A | T | DD | V | D | L |
| RS/BR/15/2S | T | I | T | Q | RRG | H | - | T | T | DD | V | D | L |
| HLJ1506-1 | T | I | T | Q | RRG | H | - | T | T | DD | V | D | L |
| 915F06007FD | N | I | T | R | GKR | Y | - | A | T | EE | A | D | L |
| G17 | T | I | T | R | RRG | Y | S | A | S | EE | A | D | L |
| CL33 | N | I | T | R | GKR | Y | - | A | T | EE | A | D | L |
| S53/It | N | I | T | R | GKR | Y | - | A | T | EE | A | D | L |
| HLJ1510 | N | I | T | R | GKR | Y | - | A | T | EE | A | D | L |

ORF2 encodes VP3, which is a non-structural protein comprising 124 amino acid residues. HB2018S1 VP3 had 92.0%–99.2% homology with the other strains. Compared with the amino acid substitution sites of the reference strains, HB2018S1 had only amino acid substitutions at site 83H/Y, excluding G17, which had an R insertion at site 122. The other major amino acid mutation sites were as follows: 7R/H, 9R/Q, 12T/I, 14Q/R, 28S/C, 54Y/S, 65A/V, 69D/A, 71G/E, 79V/A, 81L/S, 85R/K,99A/S,101K/R, 103Q/R, 104Q/R, 115N/E, 20K/R, and 124L/V (displayed in gray in Table 4).

**Table 4.** Mutations at different amino acid sites in AGV2 VP3

| Strains | 7<br>R/H | 9<br>R/Q | 12<br>T/I | 14<br>Q/R | 28<br>S/C | 54<br>Y/S | 65<br>A/V | 69<br>D/A | 71<br>G/E | 79<br>V/A | 81<br>L/S | 83<br>H/Y | 85<br>R/K | 99<br>A/S | 101<br>K/R | 103<br>Q/R | 104<br>Q/R | 115<br>N/E | 120<br>K/R | 122<br>-- | 124<br>L/V |
| --- | --- | --- | --- | --- | --- | --- | --- | --- | --- | --- | --- | --- | --- | --- | --- | --- | --- | --- | --- | --- | --- |
| HB2018S1 | R | R | T | Q | S | Y | A | D | G | V | L | Y | R | A | K | Q | Q | N | K | - | L |
| NX1510 | R | R | T | Q | S | Y | A | D | G | V | L | H | R | A | K | Q | Q | N | K | - | L |
| NX1506-2 | R | R | I | Q | S | Y | A | D | G | V | L | H | R | A | K | Q | Q | N | K | - | L |
| NX1506-1 | R | R | T | Q | S | Y | A | D | G | V | L | H | R | A | K | Q | Q | N | K | - | L |
| JL1508 | R | R | T | Q | S | Y | A | D | G | V | L | H | R | A | K | Q | Q | N | K | - | L |
| LN1511 | R | R | T | Q | S | Y | A | D | G | V | L | H | R | A | K | Q | Q | N | K | - | L |
| JX1602 | R | R | T | Q | S | Y | A | D | G | V | L | H | R | A | K | Q | Q | N | K | - | L |
| GZ1601 | R | Q | T | R | C | Y | A | D | G | A | S | H | R | S | K | R | R | E | K | - | L |
| HLJ1603-2 | R | Q | T | R | C | Y | A | D | G | A | S | H | R | S | K | R | R | E | K | - | L |
| HLJ1603-1 | R | Q | T | R | C | Y | A | D | G | A | S | H | R | S | K | R | R | E | K | - | L |
| HE1511 | R | Q | T | R | C | Y | A | D | G | A | S | H | R | S | K | R | R | E | K | - | L |
| HLJ1506-2 | R | R | T | Q | S | Y | A | D | G | V | L | H | R | A | K | Q | Q | N | K | - | L |
| GS1512 | R | R | T | Q | S | Y | A | D | G | V | L | H | R | A | K | Q | Q | N | K | - | L |
| HLJ1508 | R | R | T | Q | C | S | V | A | E | V | L | H | R | S | R | Q | R | E | K | - | L |
| G13 | R | R | T | Q | S | Y | A | D | G | V | L | H | R | A | K | Q | Q | N | K | - | L |
| JL1511 | R | R | T | Q | C | S | V | A | E | V | L | H | R | S | R | Q | R | E | K | - | L |
| HM590588 | R | Q | T | R | C | Y | A | D | G | A | S | H | R | S | K | Q | R | E | K | - | L |
| JQ690763 | R | Q | T | R | C | Y | A | D | G | V | L | H | R | S | K | Q | R | E | K | - | L |
| RS/BR/15/2S | R | Q | T | R | C | Y | A | D | G | A | S | H | R | S | K | Q | R | E | K | - | L |
| HLJ1506-1 | R | Q | T | R | C | Y | A | D | G | A | S | H | R | S | K | Q | R | E | K | - | L |
| 915F06007FD | R | R | T | Q | S | Y | A | D | G | V | L | H | R | A | K | Q | Q | N | K | - | L |
| G17 | H | Q | T | R | S | Y | A | D | G | V | L | H | K | A | K | Q | R | E | R | R | V |
| CL33 | R | R | T | Q | S | Y | A | D | G | V | L | H | R | A | K | Q | Q | N | K | - | L |
| S53/It | R | R | T | Q | S | Y | A | D | G | V | L | H | R | A | K | Q | Q | N | K | - | L |
| HLJ1510 | R | R | T | Q | S | Y | A | D | G | V | L | H | R | A | K | Q | Q | N | K | - | L |

The 5′-untranslated regions (UTRs) of HB2018S1 were between sites 224 and 351, which comprised six closely matching direct repetition (DR) regions with lengths of 22 nt. Among these, five DRs were identical (5′-GTACAGGGGGGTACGTCACCAT-3′). The different DR was the last DR. The three bases at its 3′ end were AGC of HB2018S1, which was similar with the reference strains.

## DISCUSSION

To the best of our knowledge, AGV2 has been detected in three different hosts: chickens, humans, and ferrets. Most strains of AGV2 are mainly detected in chickens, which is its original host. In the present study, AGV2 was detected, for the first time, in a cold-blooded farmed snake; however, how the snake was infected remains unknown. Farmed snakes are mainly fed on raw meats and specific feed. In this study, only one snake farm was positive for AGV2 DNA. According to the feed survey, the snakes in the positive farm were usually fed on chickens. Whether snakes are infected with AGV2 because of alimentary transmission following chicken consumption requires further investigation. In 2012, AGV2 was detected in the feather sacs of two adjacent chicken flocks in southwestern Brazil, suggesting that there is a horizontal transmission mode of AGV2 although more studies are required to demonstrate this phenomenon. From 2011 to 2013, some researchers conducted epidemiological analysis of AGV2 in serum and feces of some individuals to determine whether AGV2 could be replicated in humans. Whether the human AGV2 is attributable to metabolic residues in poultry food remains to be clarified. Phylogenetic tree analysis showed that the sequence of HB2018S1 exhibited high homology with the sequences of NX1506-1, NX1506-2, and NX1510 from China. However, the regions are far from the site of collection of HB2018S1, and the AGV2 transmission mode requires further investigation.

The AGV2 genome is 2376 nt in length, which is similar to the AGV2 genome from chickens in China, with three overlapping ORFs similar to the ones in chicken infectious anemia virus (CIAV) encoding VP1, VP2, and VP3. At the protein/amino acid level, the homology of VP1, VP2, and VP3 proteins in AGV2 corresponding with VP1, VP2, and VP3 proteins in CIAV was 38.8%, 40.3%, and 32.2%, respectively [1]. 5′-UTR, which is located between the polyadenylation site and the transcription beginning, is similar to the structure of CIAV [11, 12].

The VP1 protein of CIAV is the only capsid protein of the virus, which constitutes the neutral antigen site of the virus. There are three conserved replication motifs on the VP1 sequence in CIAV, including FATLT (at 313–317 residues), QRWHTLV (at 351–357 residues), and YALK (at 402–405 residues)[13]. Similar sequences were observed in the HB2018S1 genome: FAALS (325–329 residues), RRWLTLV (363–369 residues), and KAMA (412–415 residues). These sites are completely conserved in the currently reported AGV2 sequences. Based on a part of the AGV2 sequences, Yao et al. (2016) hypothesized that there was a high-variant region in the middle and back of the VP1 protein, ranging from 288 to 314 residues. In the present study, only eight strains of viruses, including GS1512, HLJ1508, G13, JL1511 HM590588, JQ690763, RS/BR/15/25, and G17, had mutations, and the mutation sites were not identical. Because of the limited number of sequences, it was not possible to predict the high variation regions accurately. However, further investigations are required to verify whether the five mutations in the VP1 protein in HB2018S1, which are different from the other sequences, affect the host selectivity of the virus.

The VP2 protein in CIAV is a dual-specific protein phosphatase [14]. The WX7HX3CXCX5H sequence of the phosphatase motif is also highly conserved in CIAV, TTV, and TTV-like TLMV [15]. A similar motif in the VP2 protein of AGV2 is WLRQCARSHDEICTCGRWRSH (95–115 residue) [14] performed a site-directed mutagenesis of VP2 and observed that it affected the replication of viral particles, resulted in cytopathological changes, and downregulated the levels of MHC I in infected cells. VP1 and VP2 were necessary for the replication of viral particles. Noteborn [16] also observed interaction between VP1 and VP2 proteins based on an immuncoprecipitation test, which further confirmed that the non-structural protein VP2 plays a scaffold role in virus particle assembly. In the VP2 protein, HB2018S1 has four different amino acid mutations and a S insertion site of G17 from ferrets. Whether such mutations alter VP2 function and, in turn, affect virus replication and whether they affect the scaffold function of the VP2 protein require further study.

The VP3 protein in CIAV is also known as apoptin. In normal cells, the VP3 protein exists in the cytoplasm. However, in cancerous cells, it exists in the nucleus and induces apoptosis [17, 18]. It has also been demonstrated that the VP3 protein in AGV2 induces specific apoptotic functions in tumor cells, while the change in molecular function caused by a mutation at site 83H/Y of VP3 protein in HB2018S1 still requires further experimental evidence.

In the present study, in addition to the AGV2 virus from chickens, humans, and ferrets, an AGV2 strain from a cold-blooded farmed snake was found, which could provide insight into the modes of transmission of AGV2 and its host diversity.

## MATERIALS AND METHODS

### Samples and virus detection

In 2018, a total of 91 samples (8 liver tissues from snakes died from bacterial infection and 83 faecal samples) from 7 separate king-ratsnakes (*Elaphe carinata*) farms in Hubei and Henan, China, were collected and tested for AGV2 by PCR using the AGV2-specific primers AGV2-F (5′-CGTGTCCGCCAGCAGAAAC-3′) and AGV2-R (5′-GGTAGAAGCCAAAGCGTCCAC-3′) as previously described [9]. The samples were washed with phosphate-buffered saline, repeatedly frozen and thawed thrice, immersed in liquid nitrogen, and ground into a homogenate. The homogenate was centrifuged at 5000 rpm for 10 min, and 0.2 mL of the supernatant was obtained for nucleic acid extraction. DNA and RNA were extracted using a DNA/RNA extraction kit (TransGen Biotech, Beijing, China) according to the manufacturer’s instructions, and the extracted DNA was stored at −20°C until use.

### Whole-genome sequencing of AGV2

Whole-genome sequencing of AGV2 was conducted using three pairs of overlapping primers designed by Yao et al (2016). including primers for the first segment (1F: 5′-ATT TCCTAGCACTCAAAAACCCATT-3′ and 1R: 5′-TCTGGGCGTGCTCAATTCTGATT-3′; from nucleotides 1960–379), second segment (2F: 5′-TCACAGCCAATCAGAATT GAGCACG-3′ and 2R: 5′-TTCTACGCGCATATCGAAATTTACC-3′; from nucleotides 349–1082), and third segment (3F: 5′-TATTCCCGGAGGGGTAAATTTCGAT-3′ and 3R: 5′-CCCCTGTCCCCGTGATGGAATGTTT-3′; from nucleotides 1046–2027); the amplified fragments were 802, 733, and 981 bp in length, respectively. The PCR amplification product mixed to obtain a total reaction volume of 20 μL contained a template DNA, reaction buffer, GC enhancer, 6 pmol of upstream/downstream primers, 0.4 mM of each dNTP (3 µL), and PrimeSTAR HS DNA polymerase (TaKaRa Biotechnology Co., Ltd., Dalian, China). Sequence amplification was performed under the following cycling conditions: initial denaturation at 98°C for 5 min followed by 30 cycles of denaturation at 98°C for 10 s, annealing at 60°C for 15 s, and extension at 72°C for 10 s, with final extension at 72°C for 10 min. The PCR products of the three fragments were cloned into a pMD18-T easy vector (TaKaRa Biotechnology Co., Ltd., Dalian, China) for later sequencing (Syn-Biotechnology, Suzhou, China). PCR and sequencing were performed at least thrice.

### Sequence and phylogenetic analysis

SeqMan (DNASTAR, Lasergene®, Madison, Wisconsin) was used for contig-assembly of the partial sequences. The whole-genome sequences of the snake-originated strain were submitted to GenBank with the accession number MK840982 After sequence alignment of the 25 related reference sequences available at GenBank using Clustalx v1.83, a phylogenetic tree based on nucleic acids and the deduced amino acid sequences was constructed using Mega v.6.0 using the neighbor-joining method, Kimura 2-parameter model, pairwise deletion, and bootstrap analysis of 1000 replicates [19]. In addition, mutation sites based on the amino acid sequences of three open reading frames (ORFs) were compared.

## Acknowledgments

This study was supported by the National Natural Science Foundation of China (Grant no. 31870917), the Scientific and Technological Project of Henan Province (Grant nos. 182107000040 and 182102110084), the Key Scientific and Technological Project of the Education Department of Henan province (Grant no. 18A230012), Key Scientific and Technological Project of Nanyang City (Grant nos. KJGG2018144 and KJGG2018069), and the Technological Project of Nanyang normal university (Grant nos. 16134, 18046, and 2018CX014).

